# MAGOS: Discovering Subclones in Tumors Sequenced at Standard Depths

**DOI:** 10.1101/790386

**Authors:** Navid Ahmadinejad, Shayna Troftgruben, Carlo Maley, Junwen Wang, Li Liu

**Affiliations:** College of Health Solutions, Arizona State University, Tempe, AZ, 85004, USA; Biodesign Institute, Arizona State University, Tempe, AZ, 85281, USA; Department of Health Sciences Research & Center for Individualized Medicine, Mayo Clinic Arizona, Scottsdale, AZ, 85259, USA

## Abstract

Understanding intratumor heterogeneity is critical to designing personalized treatments and improving clinical outcomes of cancers. Such investigations require accurate delineation of the subclonal composition of a tumor, which to date can only be reliably inferred from deep-sequencing data (>300x depth). To enable accurate subclonal discovery in tumors sequenced at standard depths (30-50x), we develop a novel computational method that incorporates an adaptive error model into statistical decomposition of mixed populations, which corrects the mean-variance dependency of sequencing data at the subclonal level. Tested on extensive computer simulations and real-world data, this new method, named model-based adaptive grouping of subclones (MAGOS), consistently outperforms existing methods on minimum sequencing depth, decomposition accuracy and computation efficiency. MAGOS supports subclone analysis using single nucleotide variants and copy number variants from one or more samples of an individual tumor. Applications of MAGOS to whole-exome sequencing data of 331 liver cancer samples discovered a significant association between subclonal diversity and patient overall survival. MAGOS is freely available as an R package at github (https://github.com/liliulab/magos).

## INTRODUCTION

The development of a tumor is an evolutionary process that typically initiates from a single clone and grows into a diverse population of cells via incessant mutations and selections (1-3). As a tumor progresses over time and space, different cell populations (i.e., subclones) emerge, expand and diminish, leading to a heterogeneous malignancy with multifarious clinical presentations. Understanding of this dynamic system provides valuable knowledge to facilitate early diagnosis, effective treatment and outcome monitoring of cancers (4-9).

Whole-exome or whole-genome sequencing is a common approach to studying intratumor heterogeneity (10-12). By tracking relative abundances of genomic variants in a collection of cancerous cells, scientists aim to quantify the genetic diversity of a tumor and to reconstruct the phylogeny of subclones. While single-cell sequencing is on the rise, bulk sequencing remains as the dominant technology that interrogates an amalgam of heterogeneous cells collectively and relies on *in silico* analysis to de-convolute the mixed populations. Several computational methods have been developed for this purpose, such as SciClone, PyClone and Expands (4, 13, 14). However, because these methods require a minimum sequencing depth of 100x, they are not suited for samples sequenced at a standard 30-50x depth (15). This precludes the overwhelming majority of samples sequenced to date, including those generated by collaborative consortia, such as the TCGA Pan-Cancer Atlas (average depth = 68x). Furthermore, an independent evaluation of SciClone showed that consistent subclonal characterizations can only be achieved when the depth exceeds 300x (16). New methods capable of identifying subclones reliably at a reduced sequencing depth will help discover valuable information hidden in the myriad existing tumor genomic data that currently remain unexploited.

At the algorithmic level, the constraints of sequencing depths in current methods are at least partially due to unexplained variances in bulk sequencing data. A key assumption taken by these methods is the correspondence between variant allele frequencies (VAFs, i.e., fraction of reads containing a specific mutant allele among total reads) and cellular prevalence (i.e., fraction of cells carrying this particular mutant among all cells). Subclone discovery is then translated into a task of clustering similar VAFs (13). However, VAFs is also influenced by technical factors, such as sequencing depth. Due to randomness in sequencing procedures, the same subclone may give rise to a dispersed cluster of VAFs when sequenced at a low depth, but a tight cluster of VAFs when sequenced at a high depth (16). The spread of VAFs also correlates with cellular prevalence. VAFs of variants in a common subclone are expected to scatter more broadly than those in a rare subclone (16). Without considering these confounders, subclones reported by current methods are inevitably adulterated, especially when the sequencing depth is not high enough to create strong contrasts between subclones with similar cellular prevalence.

To detangle these technical variabilities from biological variabilities, we have developed a new method, named <u>m</u>odel-based <u>a</u>daptive <u>g</u>rouping <u>o</u>f <u>s</u>ubclones (MAGOS) that explicitly models the impact of sequencing depth and cellular prevalence on the variance of VAFs in subclone decomposition. Through extensive tests using computer simulations and real-world data, we show that MAGOS can accurately delineate subclonal structures of tumor samples sequenced at depths as low as 30x. MAGOS is also the fastest program when compared to SciClone and PyClone, showing an acceleration of 3-20 folds. We implemented MAGOS as an R package that is freely available at github (https://github.com/liliulab/magos).

## RESULTS

The purpose of MAGOS, as well as SciClone and PyClone, is to group variants that emerge and evolve together into a cluster based on similarities of VAFs. Each cluster thus corresponds to a subclonal expansion. In this context, we use VAF cluster and subclone interchangeably.

### MAGOS Algorithm

MAGOS supports subclone analysis of a tumor containing single nucleotide variants (SNVs) and copy number variants (CNVs) obtained from one or more samples. To illustrate the algorithms of MAGOS, we start with simple scenarios and gradually introduce complexities into the data model.

The simplest scenario involves a single tumor sample that has only SNVs, no CNVs and no contamination of normal cells. The task is to find clusters of SNVs with similar VAFs. Given a variant *i*, we denote the total number of reads aligned to this position as its sequencing depth *e*_*i*_, and denote the fraction of reads containing the mutant allele among all reads as its VAF *v*_*i*_∈ (0, 1). For a set of *m* variants belonging to the same subclone, we model their VAFs as random samples from a beta distribution *Beta*(*α, β*) and require the two shape parameters (*α* and *β*) to satisfy

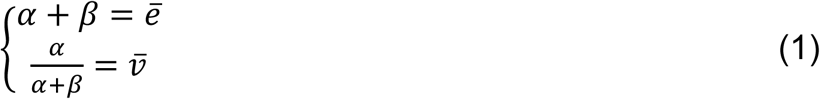

where *ē* is the mean sequencing depth and 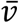 is the mean VAF of these variants. This configuration has the desired property that the variance of the beta distribution is positively correlated to the mean VAF and negatively correlated to the mean sequencing depth. Through this setup, we links the variance of VAFs to cellular prevalence and sequencing depth.

When multiple subclones are present, the observed VAFs are a mixture of samples from multiple beta distributions, each defined by a set of shape parameters. Therefore, identification of subclones is equivalent to decomposing mixed beta distributions (**Fig. 1A**). We solve this problem with a two-phase algorithm that performs agglomerative hierarchical clustering and adaptive partitioning.

**Figure 1.**
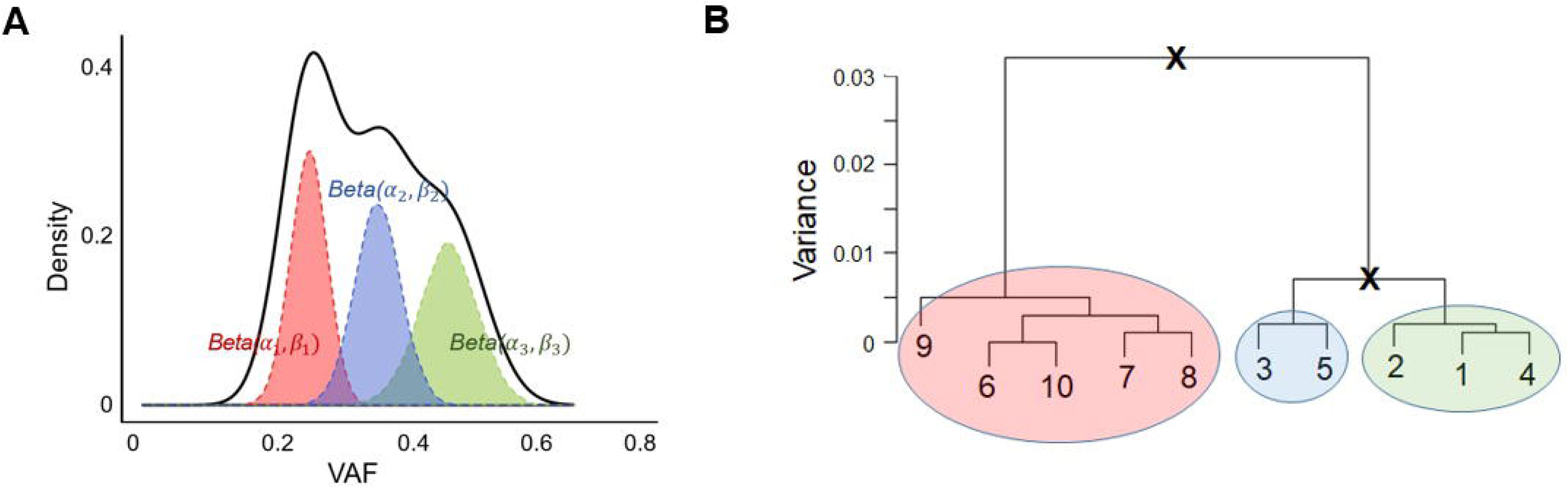
MAGOS algorithm. (**A**) Beta mixture distribution. The observed distribution (black curve) of VAFs is a combination of multiple hidden groups of VAFs, each forming a beta distribution (shaded curves) defined by different parameters. (**B**) Hierarchical clustering and adaptive partitioning. In this example, ten variants at the leaf nodes are progressively grouped into clusters based on VAF similarities to form a tree structure. To partition the tree, we follow the root-to-leave paths. At each branching point, the variance of the VAFs of the clade is compared with the expected variance. A cluster is accepted if the variance is lower than the expected value. Otherwise, it is rejected and partitioning continues (marked by black crosses). In this example, the red, blue and green cluster are accepted.

In the first phase, we organize variants into a hierarchical tree structure by progressively grouping variants with similar VAFs into a cluster. Starting with leaf nodes each consisting of an individual variant, we iteratively merge a pair of nodes with the shortest distance among all pairs to create a new cluster till all variants are merged into one root cluster. Given two nodes (i.e., clusters), *C*_1_ *and C*_2_ consisting of *m*_1_ and *m*_2_ variants, respectively, we define their distance *d* as a weighted sum of negative log likelihood that VAFs of all variants in *C*_1_ *and C*_2_ are drawn from the same beta distribution,

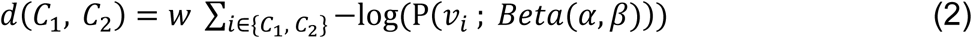

where *α* and *β* are calculated by solving equation (1) and the weight *w* = 1/(*m*_1_ + *m*_2_) · *var*(*v*) · *range*(*v*). Because the distance is down-weighted by the variance and range of VAFs, given two pairs of clusters with similar values of log likelihood, MAGOS will choose the pair with a smaller variance and a narrower range to merge at an earlier step.

In the second phase, we identify boundaries of distinct beta distributions by traversing and partitioning the tree into clades (i.e., aggregation of clusters below a branching point). Unlike traditional approaches that cut the tree at a fixed branch level, we perform an adaptive splitting (**Fig. 1B**). Along the root-to-leaves path, we examine the clade at each branching point and test the null hypothesis that VAFs in this clade are drawn from the same beta distribution. This is done by comparing the observed variance of VAFs with the expected variance of VAFs. Specifically, given a clade containing *m* variants, we assume they belong to the same subclone and compute *α* and *β* by solving equation (1). We then draw *m* random samples *x*_1:*m*_∼*Beta*(*α, β*) and calculate *var*(*x*). By repeating this process 1,000 times, we derive 1,000 *var*(*x*) values representing the null distribution. We then use one-sample one-sided t-test to evaluate if 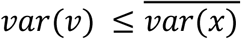. We reject the null hypothesis if the p-value < 0.01, which indicates VAFs of this clade are from heterogeneous beta distributions and needs to be partitioned further. Otherwise, we consider this clade as homogeneous and stop traversing below this branching point. We repeat this process till we find homogeneous clades along all branches or we reach the leaf nodes. Each of the resulted homogeneous clade represents a unique cluster.

When multiple samples of a tumor are analyzed, we expect that VAFs representing the same subclone change concordantly across all samples. However, because the sequencing depth and cellular prevalence of a subclone vary across samples, we need to estimate the beta distribution of this subclone in each sample separately. We then extend equation (1) to

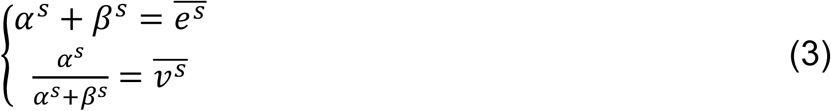

where 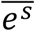 is the mean sequencing depth and 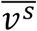 is the mean VAF of these variants in sample *s*, and *α*^*s*^and *β*^*s*^are the two shape parameters of a beta distribution specific to this sample. To determine the between-cluster distance in each tumor sample, we extend equation (2) to

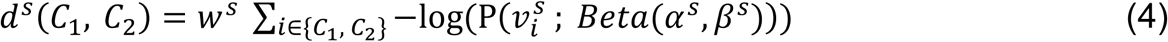

where weight *w*^*s*^ is computed for each sample *s* as *1/m* · *var*(*v*^*s*^) · *range*(*v*^*s*^). We then define the between-cluster distance across all *S* samples as

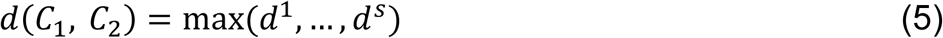

Thus, variants with concordant VAFs across all samples will be merged prior to variants with discordant VAFs. During tree partitioning, we evaluate if 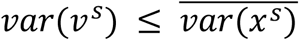 for each sample *s*. We accept a clade as a single subclone if no sample produces a p-value < 0.01.

Because the above analyses are performed on SNVs not affected by CNVs, the mean VAF 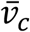 of variants in a cluster *c* is linearly correlated with the cellular prevalence 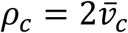 in a sample. The cluster with a mean VAF of 0.5 corresponds to heterozygous SNVs in the founding clone *f*_*c*_. If a tumor sample is contaminated with normal cells, the mean VAF of the founding clone 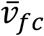 will deviate from 0.5 and the difference is proportional to the fraction of normal cells in the admixture. We then calculate the tumor purity 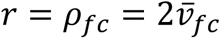 where 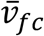 is the mean VAF of variants in the cluster closest to 0.5. Other clusters with lower VAFs consist of SNVs emerged at various time points after the founding clone expansion.

For variants located in CNV regions, we assume they do not form new clusters but instead belong to clusters identified from the SNV analysis. Given a variant *i*, the expected VAF 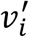 reflects the cellular prevalence *ρ*, average ploidy *φ* of the genomic region it resides and the number *k* of copies carrying the mutant allele,

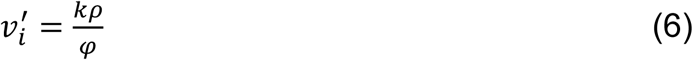

Note that *φ* is the average ploidy of the focus region in the entire sample and takes a continuous value. Finding the cluster assignment *g* from among existing SNV clusters {1, …, *C*} is to solve

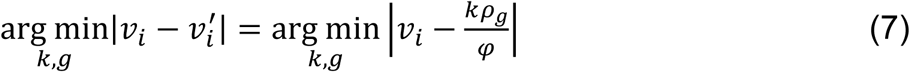

where *v*_*i*_ is the observed VAF. We limit the search space of *k* to integers between 1 and 10 × *φ*. Extensions to accommodate CNVs in multiple samples are described in Supplementary Method.

### Performance on simulated single tumor samples

To estimate the lower bound of sequencing depth and difference of mean VAFs 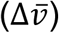 between subclones that can be detected by MAGOS and two other methods, namely PyClone and SciClone, we simulated tumor samples with a simple two-population structure. A previous study has sequenced a primary leukemia sample at various depths from 60x to over 10,000x (16). This sample had an estimated purity of 90.3% and consisted of 1,343 somatic variants in the founding clone. The distributions of VAFs of these variants confirmed that the variance of VAFs was negatively correlated with the sequencing depth (Pearson correlation coefficient= –0.54, correlation test p-value=0.02). Using these variants as a pool, we randomly drew two populations each containing 100 variants. We then adjusted the 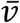 of each population between 0.05 and 0.45 with an interval of 0.05, combined these two populations and simulated read counts at an average sequencing depth of 30x, 50x, 100x, 200x, 300x and 500x via a Poisson-based down-sampling procedure (Supplementary Method). For each combination of 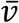 values and sequencing depth, we created 10 artificial admixtures. To quantify decomposition accuracies, we computed a weighted Jaccard index (*J*) that considered both the number of clusters identified and the assignment of variants (Supplementary Methods). A *J* score takes a value between 0 and 100, with 100 indicating a perfect match between true compositions and inferred compositions.

We first examined admixtures in which the 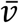 of one population was 0.45 representing a founding clone and the 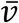 of the other population was lower than 0.45 representing a derived clone. We presented three examples to illustrate different decomposition results of these methods. In an admixture with a 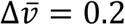 sequenced at a 30x depth, the distributions of VAFs of the two populations showed substantial overlaps covering a continuous spectrum of VAFs from 0.09 to 0.85 (**Fig. 2A**). While MAGOS found the correct number of clusters, PyClone and SciClone reported excessive clusters, mistaking the large technical variance as variance caused by mixture of multiple populations. At the 300x depth, the same admixture produced well-separated distributions of VAFs (**Fig. 2B**). Both MAGOS and SciClone then found two clusters correctly. PyClone however still reported more than two clusters with many clusters containing one or two variants. When we reduced the 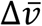 to 0.05, it was extremely challenging to divide the two populations even at the 300x coverage (**Fig. 2C**). In this case, SciClone reported one cluster and PyClone reported four clusters, respectively. Although MAGOS successfully recognized the existence of two clusters, it assigned only 75% of the variants to the correct cluster.

**Figure 2.**
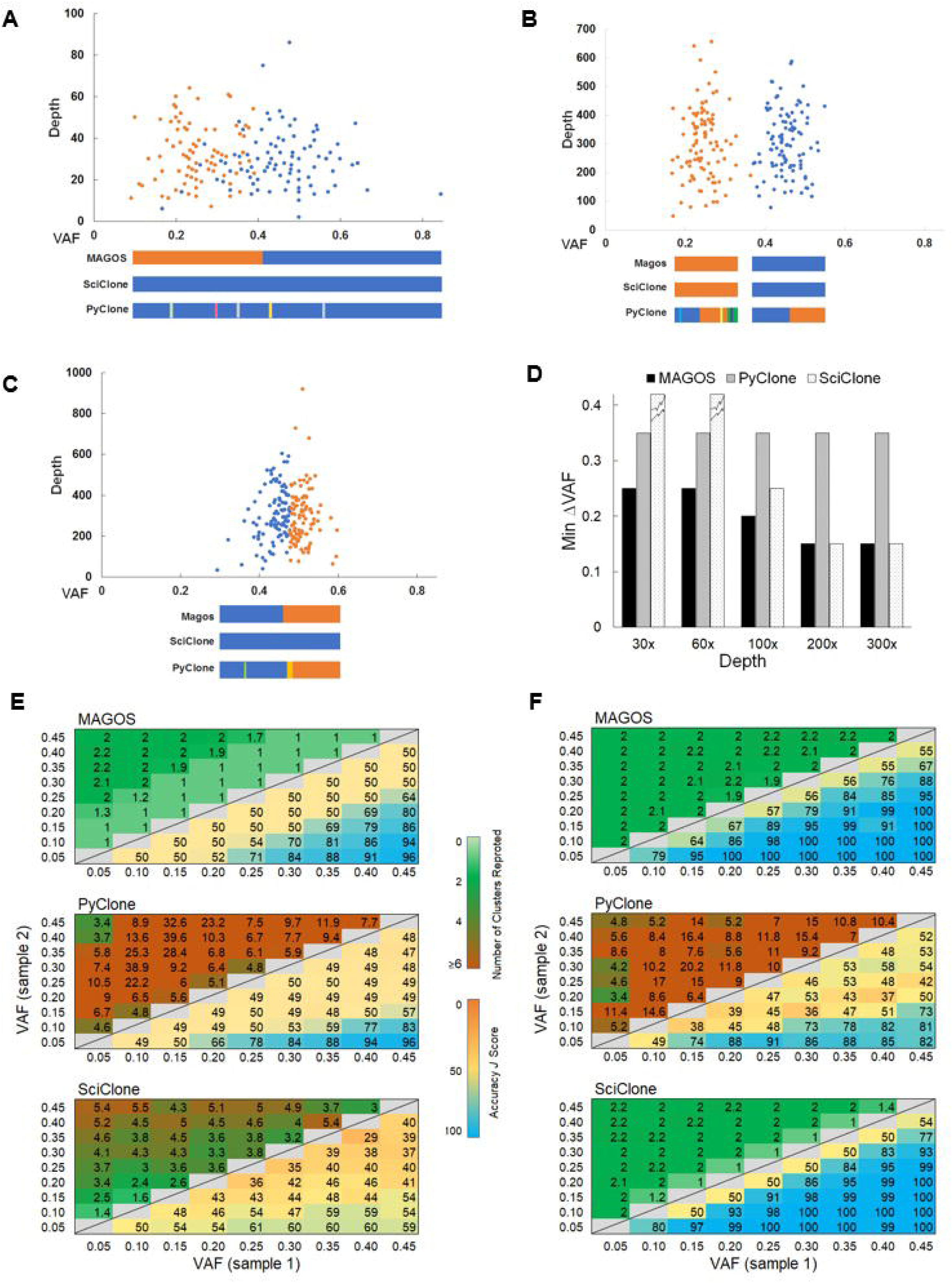
Performance of MAGOS, PyClone and SciClone on simulated single tumor samples, each consisting of two subclones. (**A, B, C**) The scatter plots show two simulated clusters of variants in a tumor sample. We varied the mean VAF of each cluster and the average sequencing depth. In panels A and B, the mean VAFs of the two clusters are 0.45 and 0.20, respectively, sequenced at an average depth of 30x (A) and 300x (B). In panel C, the mean VAFs of the two clusters are 0.45 and 0.4, respectively, sequenced at an average depth of 300x. (**D**) Minimum ΔVAF of two subclones that can be decomposed with an accuracy *J* score >80 by each method. Broken tops on bars indicate *J* scores >80 cannot be achieved. (**E, F**) Number of reported clusters (upper triangle) and *J* scores (lower triangle) at sequencing depths of 30x (D) and 300x (E). Displayed values are averages of 10 simulations. Perfect decompositions shall report 2 clusters and a *J* score value of 100.

Using *J* >80 as the accuracy threshold, we recorded the minimum 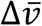 value between the two populations at a given sequencing depth for each method. The advantage of MAGOS was the most prominent at the depths of 30x – 50x (**Fig. 2D**). In these simulations, MAGOS could produce accurate decompositions with 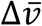 as low as 0.25. PyClone required a 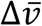 of at least 0.35. SciClone could not achieve *J* >80 at any level of 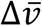. MAGOS retained the leading position till the sequencing depth increased to 200x, beyond which both MAGOS and SciClone could decompose the admixtures equally well. Interestingly, the minimum 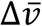 for PyClone remained at 0.35 across all sequencing depths.

Next, we examined the decomposition accuracies of all admixtures. The *J* score of all three methods was positively correlated with the 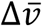 value (linear regression coefficients for MAGOS, PyClone and SciClone are 1.30, 1.03 and 0.87, respectively, all p-values <10^−12^). The *J* score was positively correlated with the sequencing depth for MAGOS and SciClone (coefficients are 0.08, 0.15, respectively, p-value <10^−16^), but not for PyClone (coefficient=0.006, p-value=0.51). At the 30x depth, MAGOS could achieve an average *J* score ≥ 80 when 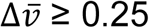 (**Fig. 2E**). In a total of 100 such admixtures, the average *J* score of MAGOS was 86.6, which was significantly better than that of PyClone (73.7, t test p-value=0.008) and SciClone (54.4, p-value=3.5×10^−8^). As the depth increased to 300x, MAGOS could achieve an average *J* score ≥ 80 when 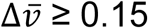 (**Fig. 2F**). In a total of 210 such admixtures, the average *J* score of MAGOS was 97.2, which was significantly better than that of PyClone (64.9, t test p-value=2 × 10^−8^) but slightly worse than SciClone (98.7, p-value=0.02).

### Performance on simulated multiple tumor samples

To evaluate the performance on delineating complicated subclonal structure embedded in multiple samples from an individual tumor, we used an established method (17) to simulate the admixtures. Each simulation contains 200 variants distributed among 3 subclones, and 40 replicated were generated at sequencing depth of 30x, 50x, 100x and 300x. The largest improvement was at 30x depth where MAGOS had a mean *J* score of 0.82 whereas SciClone and PyClone had a mean J score of 0.66 and 0.78, respectively (paired t test p-values<10^−4^, **Fig 3**). MAGOS remained at the leading performance at the sequencing depths increased to 50x and 100x (p<0.05). As the sequencing depth reached 300x, SciClone a performed equally well as MAGOS, achieving similar mean *J* scores of 0.95. Interestingly, although all three methods showed better performances at higher sequencing depth, PyClone was the least affected.

**Figure 3.**
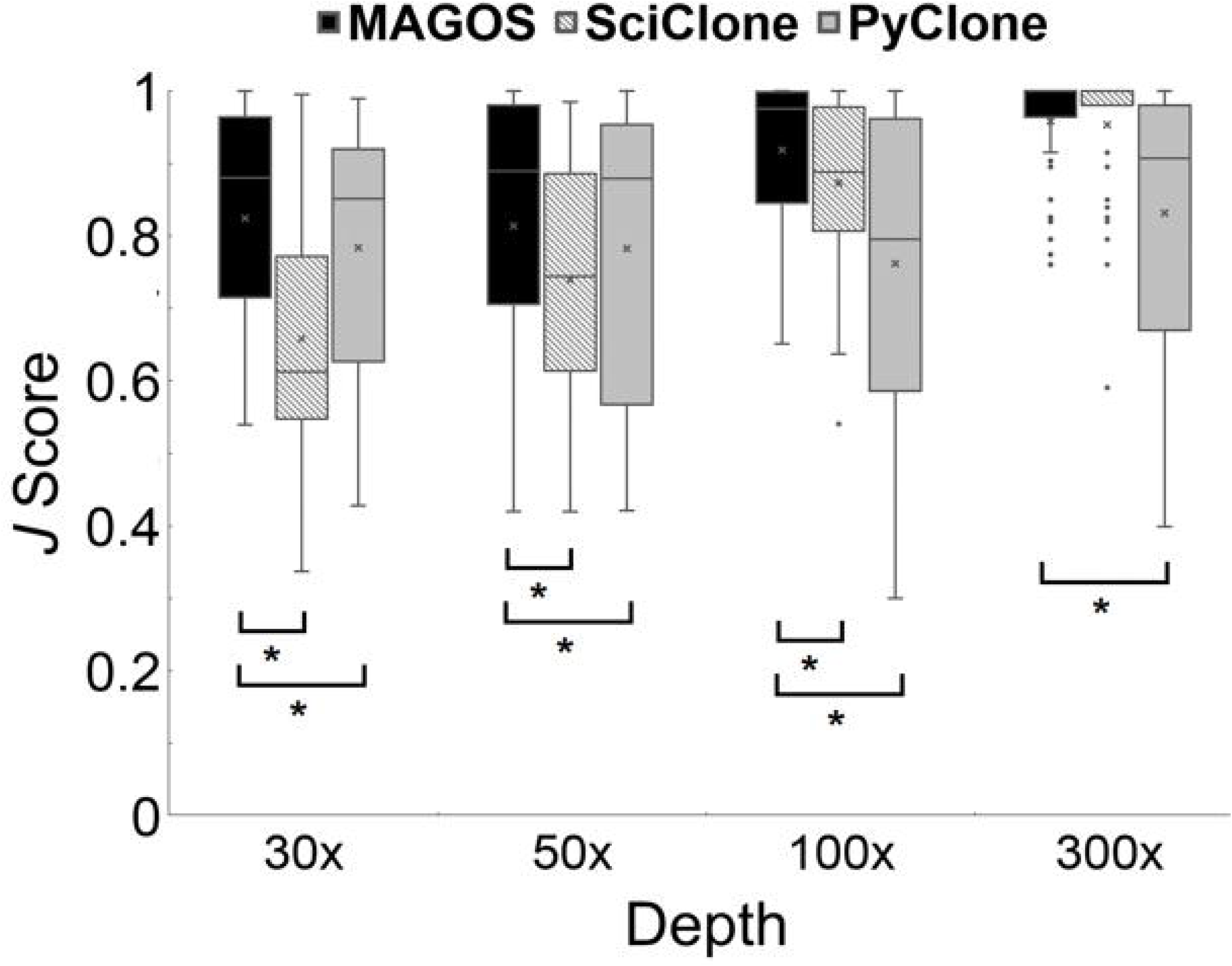
Performance on simulated multiple tumor samples, each consisting of three subclones. Accuracies of different methods tested on tumors sequenced at depth from 30x to 1,000x. Asterisks indicate a significant better performance of MAGOS as compared to the other two methods.

### Performance on real-world sequencing data

We used the ultra-deep sequencing data published by Griffith et.al (16) to assess the accuracy and reproducibility of MAGOS and the other two methods. This dataset contains 1,337 high-quality somatic SNVs detected in a primary sample and a relapsed sample from a patient with acute myeloid leukemia. Each sample was sequenced at up to 10,000x depth for validation that represents the most comprehensively sequenced tumor. Although the true subclonal structures of these samples are unknown, we followed the authors’ suggestion and used clusters detected at highest depths as the benchmark to evaluate clusters found at lower depths (data at 30x, 60x and 300x were available). The “best truth” consisted of five clusters with high confidences and two clusters with low confidences.

At the depth of 300x, MAGOS, PyClone and SciClone performed equally well, each reporting five to seven tight clusters (**Fig. 4A-C**). When the depth drops to lower than 60x, VAFs of these subclones showed large overlaps. However, MAGOS was still able to decompose the structure correctly, reporting six clusters (**Fig. 4D**). SciClone had great difficulties in separating overlapping clusters and reported only 3 subclones (**Fig. 4E**). Results from PyClone was similar to MAGOS but included small interspersed clusters (**Fig. 4F**). At the depth of 30x, MAGOS was the only method reporting the correct number of clusters and assigning variants to the correct cluster with high accuracies (**Fig. 4G**). SciClone reported results similar to 60x (**Fig. 4H**). PyClone added more than 10 small interspersed clusters in its result **(Fig. 4I**).

**Figure 4.**
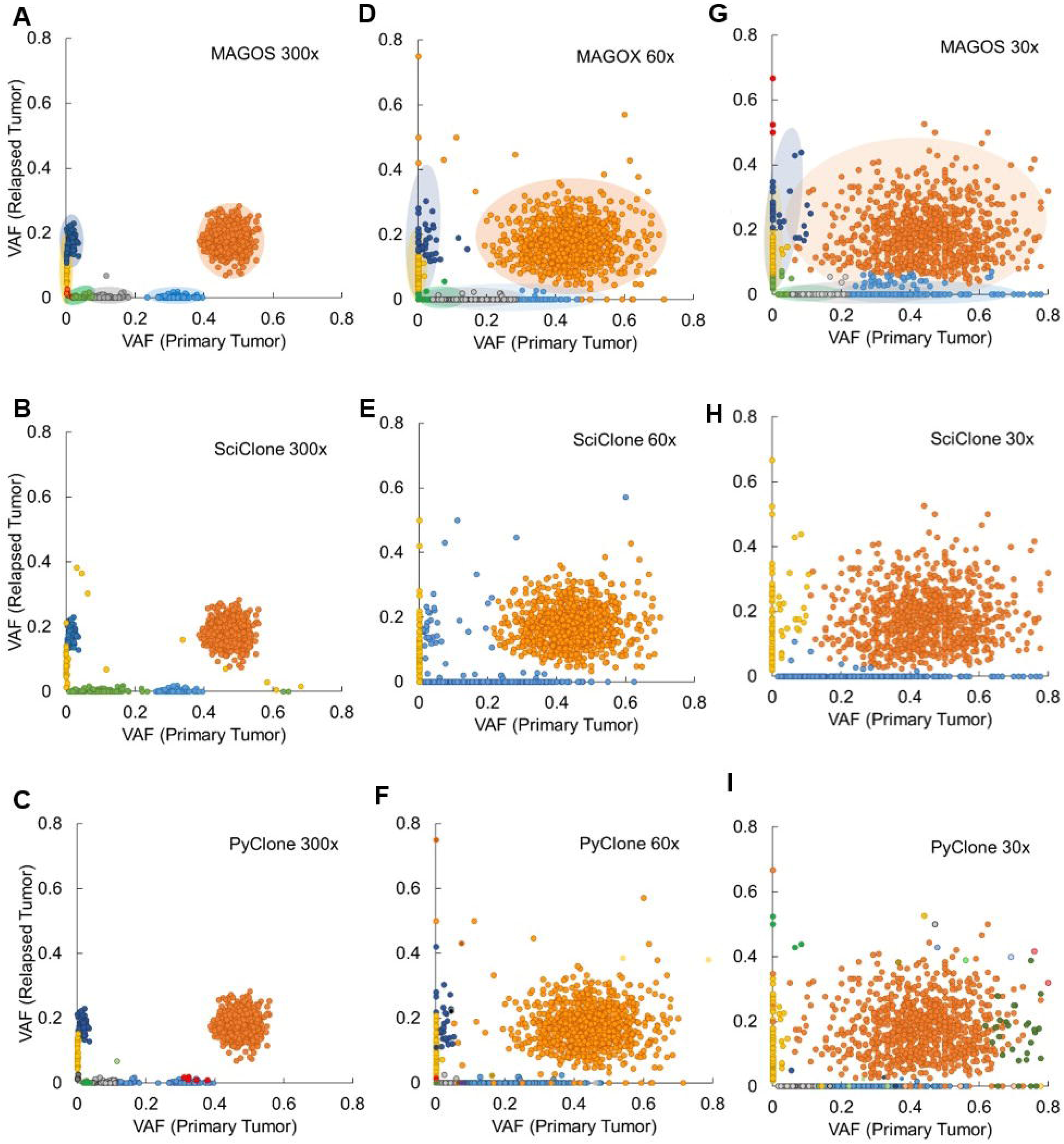
Performance on empirical data of two samples (primary tumor and relapsed tumor) from the same patient. Scatter plots show VAFs of variants sequenced at average depths of 300x (**A-C**) and 30x (**D-F**). Dots of the same color are variants assigned to the same cluster by each method. Shaded ellipses represent “true” clusters inferred from data at ∼10,000x sequencing depth. Radiuses of an ellipse correspond to 2 standard deviations of VAFs of variants belonging to a true cluster.

### Analyzing TCGA liver cancer data

We applied the MAGOS method to whole-exome sequencing data of 331 liver hepatocellular carcinoma samples from the TCGA project. The majority (79.2%) of these tumors contained 3 or 4 subclones (**Table 1**). Using Cox proportional hazard regressions, we tested if the number of subclones in a tumor was significantly associated with patient overall survival via age at diagnosis, sex and tumor stages as covariates. We found a significant association among tumors of stage III (p-value=0.01, HR=1.67, **Fig. 5**). For comparisons, the number of mutations in a tumor is not a significant prognostic factor among these tumors (p-value=0.44). Therefore, the subclone number is a novel prognostic factor for stage-3 liver cancers that is independent of age at diagnosis, sex and total number of mutations.

**Table 1.**
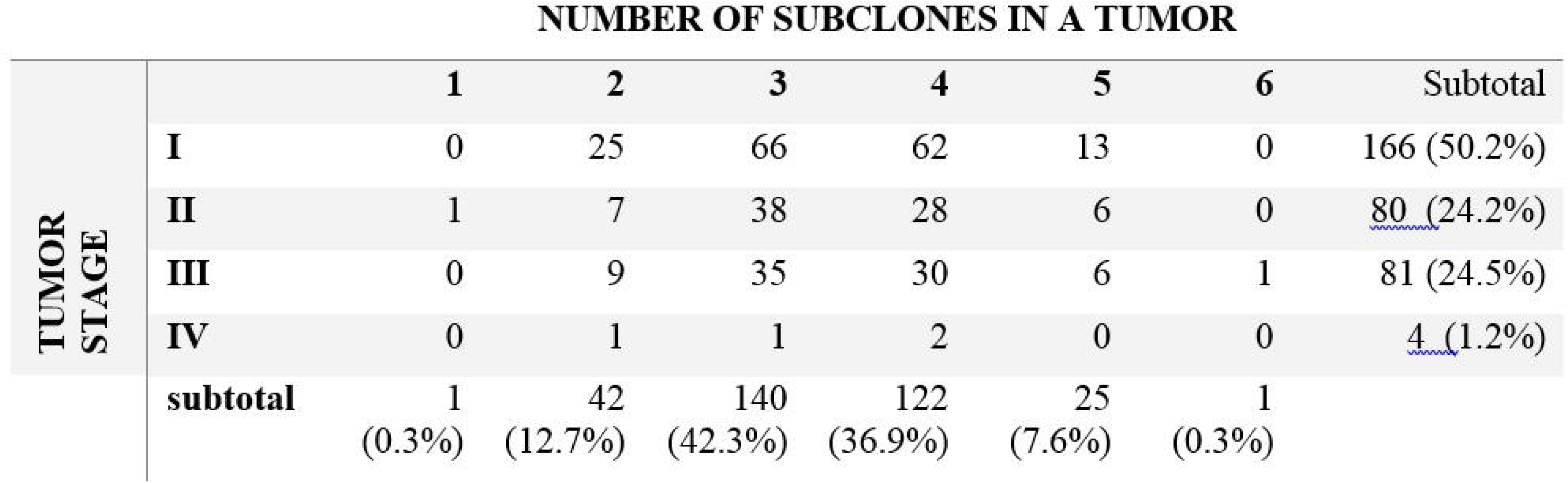

**Figure 5.**
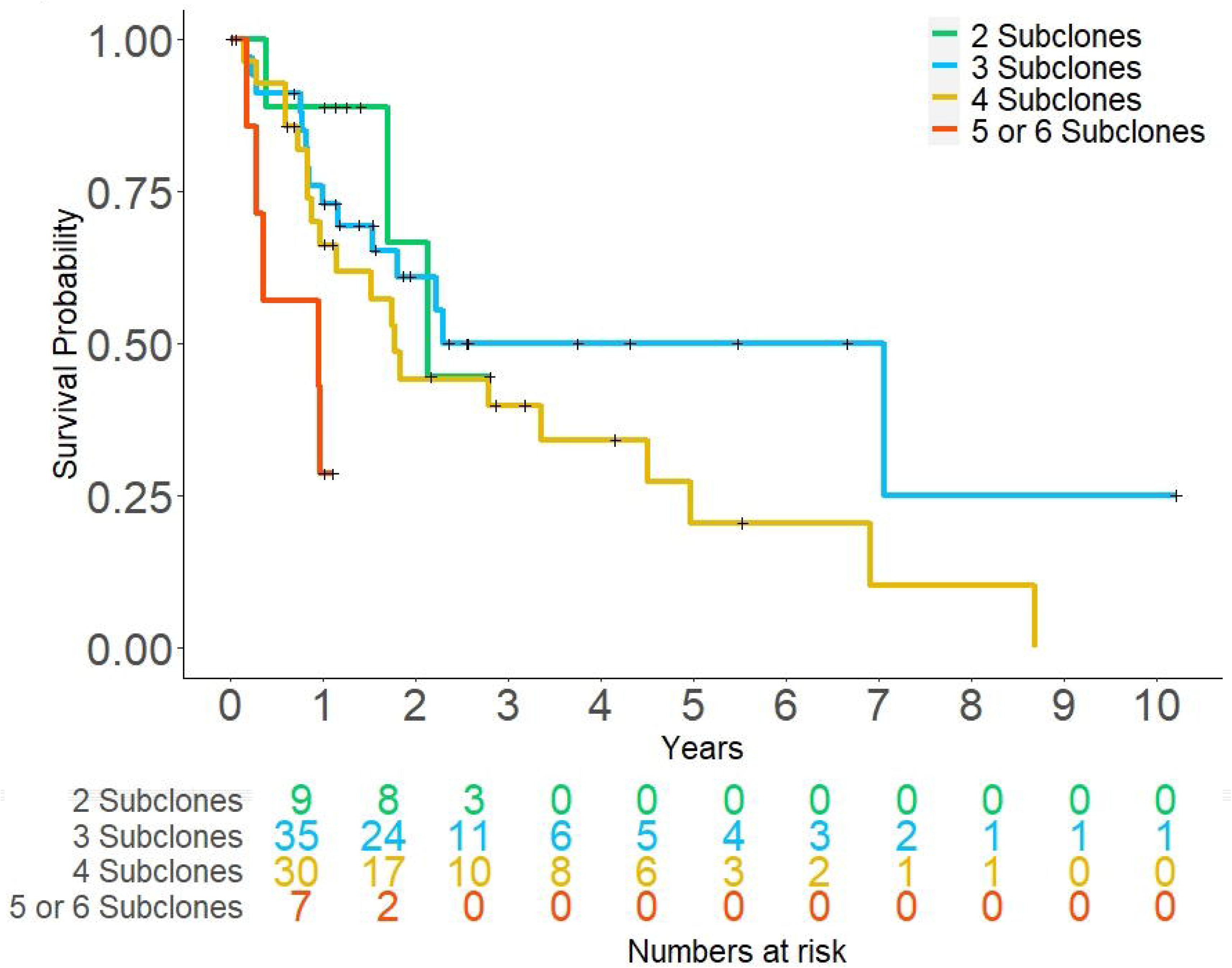
Kaplan-Meier plot of liver cancers at stage III. Tumors were stratified into groups based on the number of subclones.

## DISCUSSIONS

Cancer, as an evolutionary process, is born with a heterogeneous and dynamic nature(2, 3, 8). Precision identification and intervention of cancers shall consider the past, present and future of each tumor. With NextGen sequencing technologies, we can now catch snapshots of this process and potentially reconstruct the evolutionary history and trajectory of a tumor(6, 11, 12, 18-20). While single-cell sequencing is a promising technology to examine the genetic compositions of individual cells, uneven genome coverage, low accuracy of variant calls and prohibitive cost limit its usage in subclonal investigations (21-24). The majority of current studies and likely many others in the near future rely on bulk sequencing of mixed tumor cells and computational decomposition to identify variants that occur and evolve together. Several challenges emerge in these analyses.

First, sequencing depth is a key factor affecting the accuracy of identified subclones (15). Shown in both the simulated and real-world data (**Fig. 2-4**), as the sequencing depth drops, the centroid of each cluster remains unchanged while each cluster becomes more scattered, eventually leading to overlaps. Explicit modeling of this correlation enables MAGOS to accommodate large variances at a lower depth. However, variants in overlapping areas are impossible to separate. Instead of using VAF cutoffs to create artificial borders, a more informative measure is the probability of each variant belonging to a specific clone, which MAGOS reports.

Second, the difference of mean VAFs between subclones limits the power of distinguishing them. Increasing sequencing depth does not change the centroid of each cluster, thus helps little on detecting subclones with similar VAFs. Contrarily, additional samples from the same tumor helps segregate these clusters that are otherwise undiscernible. As MAGOS enables subclonal identifications from genomes sequenced at standard depths, the saved cost can be better invested on analyzing more samples. The benefit of sequencing additional samples is more evident at low sequencing depth, as shown in our test of PyClone. PyClone performs significantly worse than MAGOS when only a single sample is analyzed at low depth (**Fig. 2**). However, when analyzing 2 samples at depth 60x, PyClone results are similar to that from MAGOS (**Fig. 4**).

Third, as multiple samples from a tumor help identify segregating subclones and whole-genome sequencing reveals noncoding variants, these additional data also increase the computational complexity, which in turn requests efficient algorithms. We optimized the efficiency of MAGOS that tested its CPU time using simulated tumors. We varied the number of samples for each tumor, the number of mutations in each sample and the mean sequencing depth. Across all configurations, MAGOS showed 3-20x acceleration as compared to SciClone and PyClone (**Fig. 6A-C**), making it a fast and reliable method for subclone decompositions. The identified clusters can be used for further analyses, such as tumor phylogenetic inferences.

**Figure 6.**
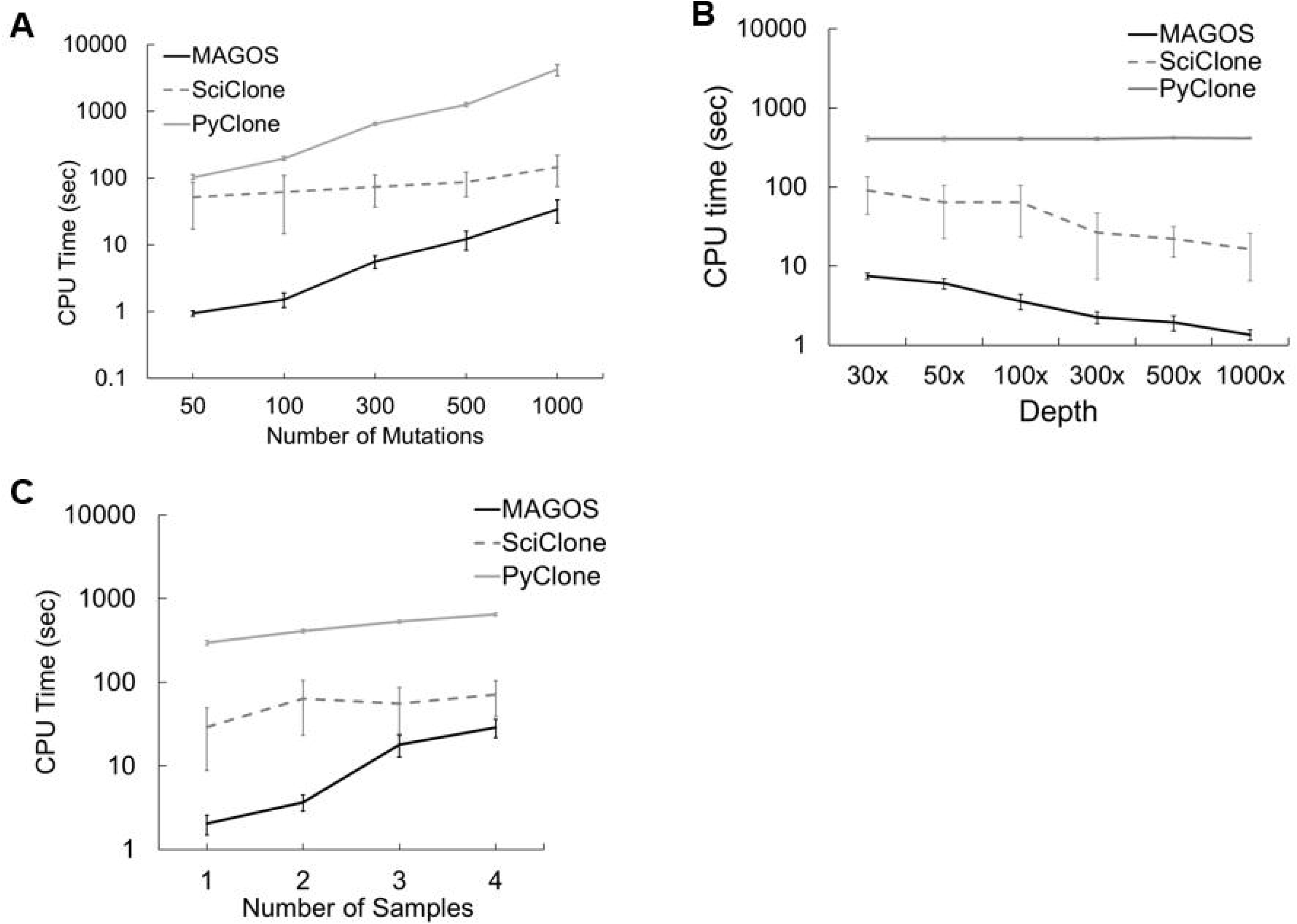
Computational efficiency comparisons tested on simulated tumors. (A) Each tumor has two samples and each sample contains 50 to 1,000 mutations sequenced at 100x depth. (B) Each tumor has two samples and each sample contains 100 mutations sequenced at depth from 30x to 1,000x. (C) Each tumor has 1 to 4 samples and each sample contains 100 mutations sequenced at 100x depth.

We implemented MAGOS as an open-source R package that is available through github (https://github.com/liliulab/magos).

