## Supplementary Methods for "MAGOS: Discovering Subclones in Tumors Sequenced at Standard Depths"

*MAGOS algorithm*

MAGOS takes two matrices ($R$ and $C$) as inputs. The matrix $R$ contains the number of reads mapped to the reference allele ($e_{r,i}^{s}$) and the number of reads mapped to the alternative allele ($e_{a,i}^{s}$) for each SNV $i$ in each sample $s$. The matrix $C$ contains the average ploidy ($\varphi_{j}^{s}$) of each CNV $j$ in each sample $s$. We categorize SNVs into two groups based on if they are located in regions affected by CNVs.

For a SNV $i$ not affected by CNVs in any sample, its sequencing depth in sample $s$ is $e_{i}^{s} = e_{r,i}^{s}+e_{a,i}^{s}$ and VAF is $v_{i}^{s} = e_{a,i}^{s}/e_{i}^{s}$. Given a collection of such SNVs, our task is to organize them into groups so that SNVs in the same group have similar VAFs that are significantly different from variants in a different group. We achieve this task with a two-phase hierarchical clustering and adaptive partitioning algorithm.

In the hierarchical clustering phase, we start with leaf nodes each consisting of an individual SNV, and iteratively merge a pair of nodes with the minimum distance among all pairs to create a new cluster till all SNVs are merged into one root cluster. We define the distance between two nodes (i.e., clusters) $C_1$ and $C_2$ as

$$d(C_1,\ C_2) = \max_{s}\left(\frac{1}{(|C_1|+|C_1|)\cdot var(v^s)\cdot range(v^s)}\sum_{i\in\{C_1,\ C_2\}} -\log(\mathrm{P}(v_i^s\ ;\ Beta(\alpha^s,\beta^s))\right) \qquad [O1]$$

where $v^s = \{v_i^s\}$ for $i \in \{C_1,\ C_2\}$, and $\alpha^s$ and $\beta^s$ are the two shape parameters of a beta distribution computed by solving

$$\begin{cases} \alpha^s + \beta^s = \overline{e^s} \\ \frac{\alpha^s}{\alpha^s+\beta^s} = \overline{v^s} \end{cases} \qquad [O2]$$

where $e^s = \{e_i^s\}$ for $i \in \{C_1,\ C_2\}$.

In the adaptive partitioning phase, we traverse the tree along the root-to-leave path. At each branching point, we examine the clade that includes all SNVs below this point. If a clade contains $m$ variants that belong to the same subclone, VAFs of these variants in each sample must come from the same beta distribution, $v^s \sim Beta(\alpha^s,\beta^s)$. To test this hypothesis, we first calculate values of $\alpha^s$ and $\beta^s$ by solving equation [O2] for each sample $s$. We then draw $m$

random samples $x_{1:m}^s \sim Beta(\alpha^s, \beta^s)$ and calculate $var(x^s) = \sum(x^s - \overline{x^s})^2 / m$. By repeating this sampling process 1,000 times, we derive 1,000 $var(x^s)$ values representing the null distribution. Using the one-sample one-sided t-test, we assess if $var(v^s) \leq \overline{var(x^s)}$. A p-value >0.01 indicates that VAFs of this clade in sample $s$ are indeed from the same beta distribution (i.e., homogeneous). We perform this test for all samples and accept the null hypothesis if all samples produce p-values >0.01. Then, we stop traversing below this branching point and consider variants in this clade constitute a subclone. Otherwise, we will partition this node by following the path and examine the clades at the next lower level. We repeat this process till we find homogeneous clades along all branches or we reach the leaf nodes. Each of the resulted homogeneous clade then represents a unique subclone.

Next, we match SNVs located in CNV regions to subclones identified from the above analysis. Given a SNV $j$ located in a CNV region that has an unknown copy number $k^s$ carrying the mutant allele in sample $s$, we find the best match subclone $g$ by solving

$$\arg\min_{k^s, g} \sum_s \left| v_j^s - \frac{2k^s \overline{v_g^s}}{\varphi_j^s} \right|$$

where $\varphi_j^s$ is the average ploidy of the region harboring this variant, and $\overline{v_g^s}$ is the mean VAF of subclone $g$ identified from the previous analysis using only SNVs not affected by CNVs. We limit the search space of $k^s$ to integers between 1 and $10 \times \varphi_j^s$.

*Optimization to improve computational efficiency*

The standard hierarchical agglomerative clustering procedure requires calculations of all pairwise distances at each step, which leads to an exponential increase of computational complexity as the number of variants grows. However, given the narrow range of VAFs between 0 and 1, not all pairwise comparisons are necessary, especially for variants with highly similar VAFs across all samples. To find these variants, we first compute the Euclidean distance between each pair of variants based on their VAFs in all samples. We then construct an incidence matrix and set the entry in row x and column y to 1 if the Euclidean distance between variants x and y is less than 0.01. Using an undirected graph created from this incidence matrix, we search for complete graphs and collapse variants belonging to each complete graph into a leaf-node in our initial hierarchical clustering step. Because the number of leaf-nodes determines the computational complexity of a bifurcating tree, this graph-based reduction step accelerates the speed of MAGOS significantly.

*Simulating subclones in single tumor samples*

In each simulation, we create an artificial tumor sample containing two subpopulations sequenced at an average depth of $e$. Variants in each subpopulation are drawn from a pool that is the founding clone in the primary acute myeloid leukemia sample sequenced at >10,000x depth by Griffith et. al [cite]. Because this sample has an estimated purity of 90.3% and the founding clone contains only heterozygous somatic mutations, the mean VAF of variants in this pool is $u = 0.451$. To create a subclone containing $m$ variants with a mean VAF $v$, we first draw $m$ random variants from the pool. For a given variant, there are $e_r^0$ number of reads mapped to the reference allele and $e_a^0$ number of reads mapped to the alternative allele in the pool. We down-sample these reads to the lower sequencing depth $e$ according to Poisson distributions

$$\begin{cases} e_r = Pois\left(e_r^0 \eta \frac{1-v}{1-u}\right) \\ e_a = Pois\left(e_a^0 \eta \frac{v}{u}\right) \end{cases}$$

where $\eta = e/(e_r^0 + e_a^0)$, and $e_r$ and $e_a$ are the number of reads mapped to the reference allele and to the alternative allele in the simulated tumor sample, respectively. Using this strategy, we generate two subclones each containing $m = 100$ somatic variants and combine them to create an admixture, representing a single tumor sample with a two-subclone structure. We vary the mean VAF $v$ of each subclone between 0.05 and 0.45 and the overall sequencing depth $e$ at 30x, 60x, 100x, 200x, 300x, 500x.

*J* score

The *J* score is a modification of the $\%G_E$ value used by Miura et. al. to evaluate subclone discovery methods [cite Miura]. However, the *J* score accommodates subclone sizes (i.e. number of variants in each subclone) when quantifying the similarities between two sets of clusters. Specifically, we denote $T$ as a set of clusters representing the ground truth, and $D$ as a set of clusters representing predictions. For each cluster $t \in T$, we find its most similar cluster $d \in D$ based on the Jaccard Index

$$I = \frac{|A \cap B|}{|A \cup B|}$$

where $A$ is the set of variants defining cluster $t$, and $B$ is the set of variants defining cluster $d$. After all truth clusters are matched, if there remain unmatched clusters in $D$, we use the Jaccard Index to find their most similar clusters in $T$. After finding the best matched pair for all truth clusters and all predicted clusters, we compute the $J$ score as

$$J = \frac{n_p I_p}{\sum_p n_p} \times 100$$

where $I_p$ is Jaccard Index of the $p^{th}$ pair, and $n_p$ is the number of variants in the truth cluster involved in this pair. J score takes a value between 0 and 100, in which 0 means no overlap between any truth clusters and any predicted clusters, and 100 means perfect matches between truth and predictions.

*Execution of SciClone and Pyclone.*

We downloaded the SciClone program and the PyClone program from GitHub, and used default settings in the analyses.
